# Nature and biological irrelevance of mixed-type enzyme inhibition

**DOI:** 10.1101/2022.12.20.521168

**Authors:** Alessandro Pesaresi

## Abstract

Mixed-type enzyme inhibitors were originally envisaged to decrease the enzyme affinity for the substrates and the maximum turnover rate by simultaneously targeting two distinct protein sites, i.e., the active site and an allosteric site. After a century from the first formulation of this hypothesis, the consensus on its validity is still unanimous, although several of its implications are in open conflict with the current knowledge on molecular recognition mechanisms. In particular, there is no plausible explanation for the experimental evidence that mixed-type inhibitors bind the enzyme active sites always more effectively than the allotopic sites. In an attempt to solve this controversy, it was found that the preference of mixed inhibitors for active sites emerges as an inevitable numerical artifact that is implicit in the equations used to model the apparent mixed inhibition caused under certain circumstances by active site-bound competitive inhibitors. Hence, proving that the consolidated model of mixed inhibition is incorrect and, more generally, strongly pointing to the biological irrelevance of mixed-type inhibition.

## 1. Introduction

Enzymes perform all of the basic chemical tasks needed to sustain life. The control of their activity relies on a hierarchy of molecular events that range from the regulation of gene expression down to the finer activity modulation exerted by allosteric effectors and inhibitors [1–3]. Because all enzymes are subjected to inhibition, for example by substrate analogs or reaction products, enzyme inhibition constitutes the primary control mechanism of enzymatic activities [4], with implications that span through all fields of molecular biology and pharmacology [5]. Indeed, the study of enzyme inhibition by poisons, drugs and endogenous modulators has been one of the main focuses of biochemical research since the origin of enzymology.

According to the consolidated one-century old theory [6], reversible enzyme inhibition falls within two opposite extremes, namely, competitive and uncompetitive inhibition. Competitive inhibitors bind the enzyme active site in a mutually exclusive fashion with the substrate, causing a reduction in the enzyme-substrate affinity that is overcome by high substrate concentrations. Uncompetitive inhibitors, in contrast, bind sites distinct form the active site, hereinafter referred to as allotopic sites, causing a reduction of the maximal turnover rate that is substrate-independent.

Figure 1 provides the essentials of enzyme inhibition. Briefly, the fundamental parameters of enzyme kinetics, *V*_*max*_ and *K*_*M*_*/V*_*max*,_ can readily be derived by plotting the reciprocal of the initial velocities (1/V) as a function of the reciprocal of the substrate concentration (1/[S]). Competitive inhibitors, which affect *K*_*M*_/*V*_*max*_ but not *V*_*max*_, generate a bundle of lines intersecting the 1/V axis, while uncompetitive inhibitors, which affect *V*_*max*_ but not *K*_*M*_/*V*_*max*_, produce parallel lines. The inhibition constant of competitive inhibitors corresponds to the dissociation constant of the E-I complex, (*K*_*ic*_), and for uncompetitive inhibitors to the dissociation of the ternary ES-I complex, (*K*_*iu*_).

**Figure 1.**
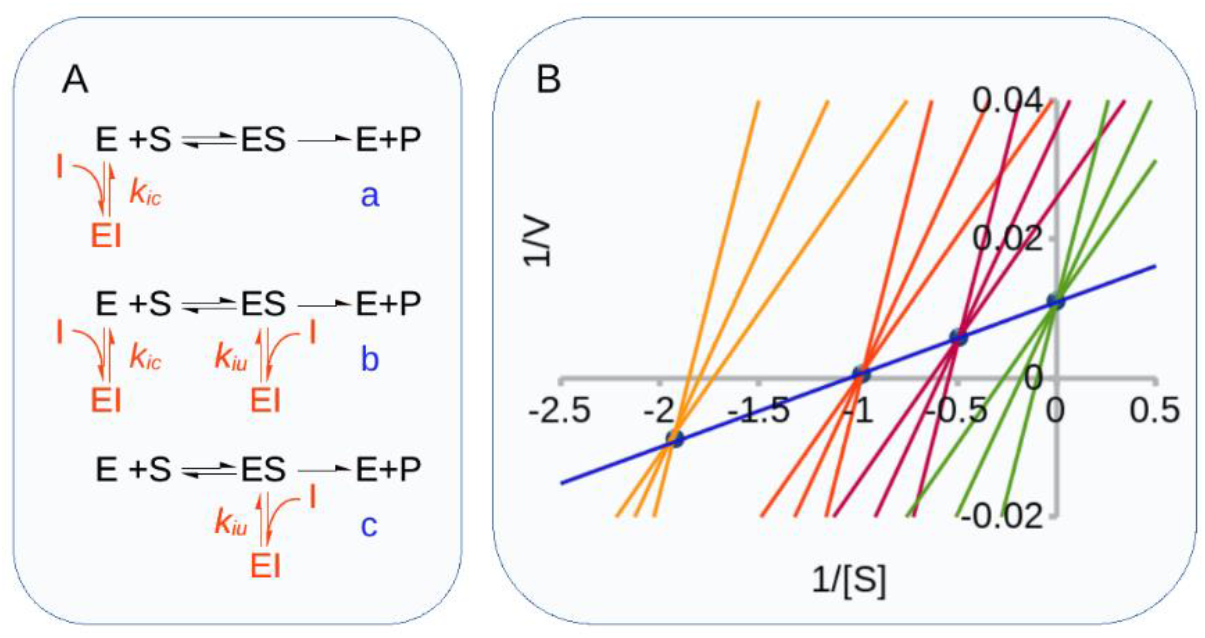
Inhibition mechanisms and patterns. **Panel A**. Inhibition mechanisms. Competitive inhibition (a), mixed-type inhibition (b), uncompetitive inhibition (c). **Panel B**. Inhibition pattern of competitive and mixed-type inhibition. In a 1/V versus 1/[S] plot of initial rates, competitive inhibitors generate a bundle of lines (green) that intercept the non-inhibited reaction trace (blue) on the 1/V axis. Mixed-type inhibitors reduce both *1/V* and *V/K*_*M*_ by binding the free enzyme (E) and the ES complex. They generate bundles of lines that intercept on the left of the 1/V axis at 1/[S] values equal to -*K*_*IC*_/*K*_*IU*_. For pure noncompetitive inhibitors (*K*_*IC*_*=K*_*IU*_, orange lines), the intercept is on the 1/[S] axis. If *K*_*IC*_<*K*_*IU*_, (red lines, *K*_*IC*_/*K*_*IU*_*=*0.5), the intercept is above the 1/[S] axis, and it is below if *K*_*IC*_>*K*_*IU*_, (yellow lines, *K*_*IC*_/*K*_*IU*_*=*2).

A third class of inhibitors, referred to as mixed competitive/uncompetitive inhibitors (or, less appropriately as noncompetitive inhibitors [7]), were originally found to decrease both *K*_*M*_/*V*_*max*_ and *V*_*max*_. By analogy with the two previous mechanisms, they were envisaged to act by targeting both the enzyme active site and the allotopic site. Therefore, the characterization of mixed-type inhibition requires the determination of a *K*_*ic*_ and *K*_*iu*_ couple. In a double reciprocal plot, mixed inhibitors generate lines that intersect at a point on the left of the 1/V axis: if *K*_*ic*_=*K*_*iu*,_ the line intersection is on the 1/[S] axis, if *K*_*ic*_<*K*_*iu*_ is above the 1/[S] axis and below if *K*_*ic*_>*K*_*iu*_ (Figure 1).

After a century from the first conceptualization by Michaelis and coworkers [6], this theory of enzyme inhibition is still accredited by unanimous consent. However, while the model of competitive inhibition is corroborated by countless evidence, for mixed-type inhibition the correlation between theory and experimental evidence remains, to a large extent, controversial. There are three main aspects in the theoretical model of mixed-type inhibition that are in open conflict with the experimental evidence and that should raise skepticism on its underlying mechanism. In order of increasing relevance these are:

i. Lack of structural evidence for the binding of mixed inhibitors to allotopic sites.
ii. Implausibility of pure noncompetitive inhibition.
iii. Mixed inhibitors always bind the active sites more effectively than allotopic sites (i.e., *K*_*iu*_≥*K*_*ic*_).

As of November 2022, in the Protein Data Bank (PDB, [8]), 118000 crystallographic structures of enzymes have been deposited. A non-exhaustive count of the enzyme-inhibitor complexes showed that more than 20,000 of these entries contain ligands defined as “inhibitors” and for which either a *K*_*i*_ or an IC_50_ is available. Because mixed inhibitors account for approximately 20% of the overall inhibition cases (see Results), the PDB should contain at least 4000 structures of enzymes in complex with mixed-type inhibitors of which, however, there seem to be no evidence. None of the 197 PDB entries that currently contain the term “noncompetitive inhibitor” or “mixed-type inhibitor”, in fact, reports ternary enzyme-substrate-inhibitor complexes or show inhibitors bound to sites other than the enzyme active site. Which constitutes a first degree of incompatibility with the two-site model of mixed-type inhibition.

Pure noncompetitive inhibition is a special case of mixed inhibition characterized by identical values of the two inhibition constants (*K*_*iu*_*=K*_*ic*_) and represents an even more puzzling phenomenon. It implies that two different molecules, the free enzyme E and the ES complex, which are different in every other respect, have identical binding constants for the inhibitor. This might be possible for very small binders, such as protons or metal ions [9], but it is extremely unlikely otherwise. Nevertheless, noncompetitive inhibition is commonly reported to occur with larger organic molecules such as drugs or metabolic inhibitors (see Results), further adding to the implausibility of the accepted model of mixed inhibition.

The last and the most striking of the three inconsistencies is that, with virtually no exception, in mixed inhibition, the uncompetitive component is always weaker than the competitive one (*K*_*iu*_≥*K*_*ic*_). This observation was first made by Cornish-Bowden on the basis of anecdotal evidence [10], and we confirmed it through an objective statistical analysis of the inhibition data stored in the BRENDA database [11], (see Results). Because there is no conceivable reason why ligands that are capable of binding both E and ES should inevitably bind E more effectively than ES, this feature of mixed inhibition seriously undermines the credibility of the accepted theory and of the published data.

Here, the hypothesis under test is that the explanation of these inconsistencies simply resides in the nonexistence of mixed inhibition, or at least, not in the form it has been theorized. An important point that needs to be addressed is that, as we are now aware, there are several exceptions to the paradigmatic rigidity of the Michaelian classification of the inhibition mechanisms. Competitive inhibition, for example, can be mimicked by allosteric effectors [12–17] and mixed-type inhibition can be caused by ligands that target only the active site. Indeed, there are five main mechanisms, or circumstances, by which an active site-bound inhibitor might appear to be mixed-type or noncompetitive when analyzed by steady-state kinetics [18–20]. Aiming to solve the awkward inconsistencies implicit in the theory of mixed-type inhibition, these five “mixed inhibition-mimicking” (MIM) mechanisms have been thoroughly reconsidered. We have derived a whole new set of equations that, surprisingly, have eluded enzymologists for over a century and that by linking *K*_*ic*_ to *K*_*iu*_ explain why, in the apparent mixed inhibition caused by each of the MIM mechanisms, *K*_*iu*_ is necessarily always larger than, or at best equal to, *K*_*ic*_.

The lack of mixed inhibition cases where *K*_*ic*_>*K*_*iu*_ suggests that the MIM mechanisms account for all, or nearly all, of the tens of thousands of inhibitors that over the years have been reported to be of the mixed or noncompetitive type. This implies that mixed-type inhibition exists mainly as a mathematical artifact, that the reported mixed inhibitors indeed only bind the active site and that the conventional 100-year-old model of mixed inhibition is inherently wrong.

## 2. Results

### 2.1. Statistics of enzyme inhibition

The quantitative statistical analysis of the available kinetic data is a precondition to turn anecdotal evidence into more solid objective observations and hence to draw conclusions on the veracity of the mixed-inhibition theory. Unfortunately, none of the extant repository stores information about the mechanism of inhibition in a structured and systematic way. In the BRENDA database [11], for example, which contains entries for more than 220,000 enzyme inhibitors and 200,000 reference citations, this kind of information has only occasionally been reported by the database curators and can be found as scattered notes in the commentary fields of a minority of the entries. Because these entry notes are not searchable, to build a reliable statistic, the relevant data were mined from the downloadable BRENDA text file. A search in this almost five million lines document of the terms “competitive”, “noncompetitive”, “mixed” and “uncompetitive” inhibitors returned a total of 7563 hits. On the basis of the occurrence of each term and in agreement with what had already been reported [4], competitive inhibition was the most common type of inhibition, accounting for 73% of total cases. Mixed/noncompetitive and uncompetitive inhibition accounted for 22% and 5% of cases, respectively. To deepen the “*K*_*ic*_ versus *K*_*iu*_ issue”, the 464 entries containing the term “mixed” or “mixed-type” inhibition/inhibitor were further analyzed. These entries point to 266 reference articles, from which it has been possible to retrieve 467 *K*_*ic*_-*K*_*iu*_ couples. If mixed inhibition was caused by the binding of inhibitors to two distinct sites, the dominance of the competitive or of the uncompetitive component would be equally probable, and a symmetric and rather flat distribution of the *K*_*ic*_/*K*_*iu*_ and *K*_*iu*_/*K*_*iu*_ ratios around a central value of 1 would be expected. The analysis of these *K*_*ic*_-*K*_*iu*_ values, however, showed that the dominance of the competitive component is overwhelmingly preponderant (*K*_*ic*_≤ *K*_*iu*_ in 90% of the cases) and that pure noncompetitive inhibition occurs with an unrealistic frequency (*K*_*ic*_/*K*_*iu*_=1 in 20%of cases), (Figure 2). It is worth noting that the large majority of the gathered *K*_*i*_s were originally obtained by the graphical analysis of initial velocity data, a method that is known to be rather inaccurate [21,22]. If to account for this inaccuracy some allowance is granted and inhibition is considered to be pure noncompetitive up to a *K*_*ic*_/*K*_*iu*_ ratio between 0.5 and 2 (rather than exactly equal to 1), then the frequency of pure noncompetitive inhibition jumps to 40%, while the occurrence of mixed inhibition cases with *K*_*ic*_*>K*_*iu*_ drops to a mere 5%. Which, evidently, is incompatible with the conventional two-site model of mixed inhibition.

**Figure 2.**
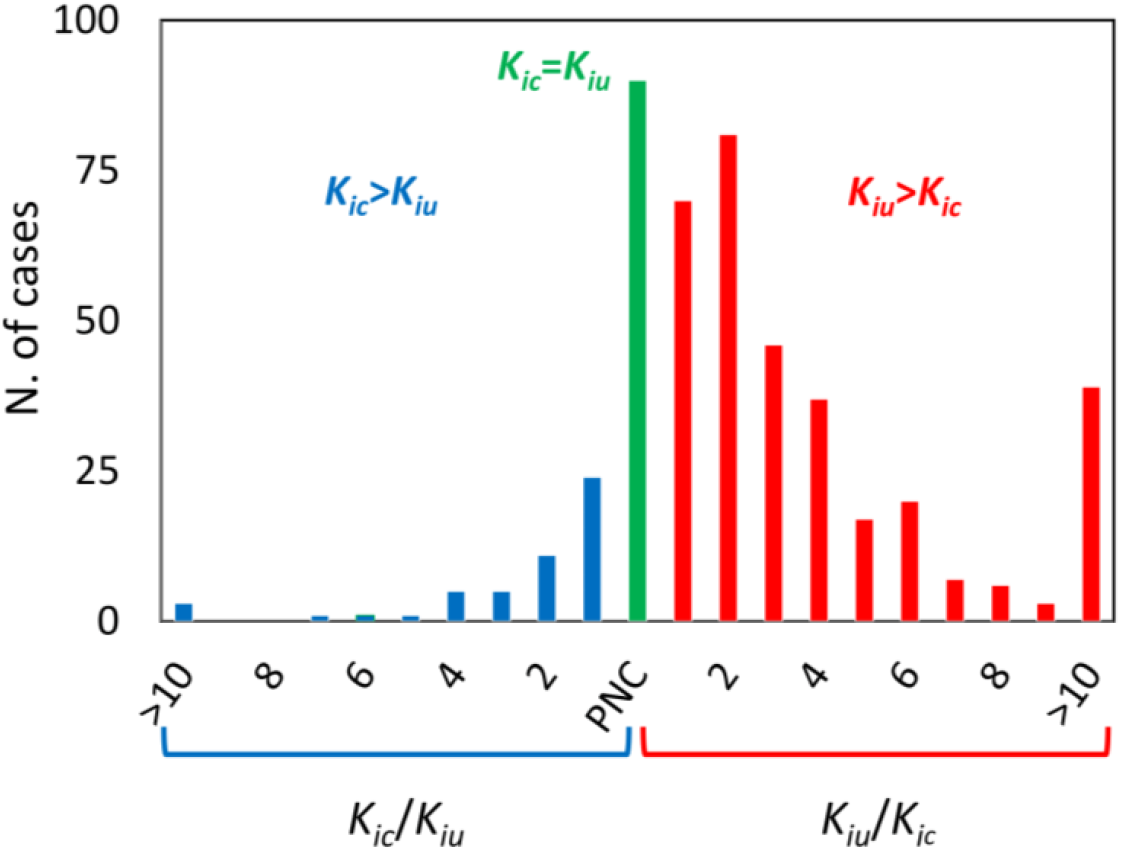
Distribution of the *K*_*iu*_-*K*_*ic*_ ratio frequencies of mixed-type inhibitors. For inhibition dominated by the competitive component (*K*_*iu*_>*K*_*ic*_, red bars), the data are presented as *K*_*iu*_/*K*_*ic*_. Data of mixed-inhibitions with *K*_*ic*_>*K*_*iu*_ (blue bars) are presented as *K*_*ic*_/*K*_*iu*_ ratio. Pure noncompetitive inhibition (PNC) is shown in green. The competitive component is prevalent in 70% of cases, pure noncompetitive inhibition represents 20% of cases, while the dominance of the uncompetitive component is observed only in 10% of cases. The total number of cases analyzed was 467.

### 2.2. Investigation into the Mixed Inhibition-mimicking Mechanisms

MIM mechanisms have been studied using KinTek Explorer [23,24], a software developed to solve enzyme kinetics by the global fitting of the reaction full time-courses through numerical integration. The program also implements a module for the simulation of enzymatic reactions once that mechanism and initial conditions are given. To determine *K*_*ic*_s and *K*_*iu*_s, as in a real experiment, the initial velocities of the progress curves computed for each MIM mechanism were analyzed by the conventional double-reciprocal plot of 1/V vs 1/[S], (see Methods).

The five MIM mechanisms have been subgrouped into two types: in the mechanism of type I, the emergence of the apparent uncompetitive component is caused by noncompliance with steady-state assumptions. In those of type II, the apparent mixed inhibition is a direct consequence of the catalytic mechanism and of the active site geometry.

#### 2.2.1. MIM mechanisms of type I – Noncompliance with steady-state assumptions

The classic steady-state analysis of enzyme kinetics is based upon two fundamental assumptions: that the substrate and inhibitor are in concentrations much higher than the enzyme and that the establishment of thermodynamic equilibrium between all the reactants settles rapidly on the steady-state time scale. With certain inhibitors, one or both of these two conditions do not apply, and consequently, the conventional steady-state treatment is inadequate.

##### Case I – Tight-binding inhibitors

To determine the *K*_*i*_ of an inhibitor, reactions must be measured in the presence of inhibitor concentrations close to the *K*_*i*_. This implies that for very potent inhibitors, typically with *K*_*i*_ in the nM range or less, the inhibitor might turn out to be not in concentration higher enough than the enzyme, hence violating the first of the two steady-state assumptions. Methods to analyze tight inhibition have been proposed [25,26], but if this circumstance is overlooked and kinetic data are analyzed according to the classic steady-state method, activesite directed inhibitors can easily be mistaken for mixed-type inhibitors [27,28].

Figure 3A shows the inhibition caused by competitive inhibitors of different potencies supplied at concentrations between 10·[E] and 0.1·[E]. In the simulation, for all inhibitors, the [I]/*K*_*i*_ ratio was set to 10. Therefore, in principle, for all of the reactions, the degree of inhibition should be the same. However, as the [I]/[E] ratio drops below 10, the inhibitor depletion that occurs upon E-I binding can no longer be neglected, and the fraction of the inactive inhibitor-bound enzyme starts to deviate from the expected linear behavior. In the double reciprocal plot, this is visualized as both a slope and an intercept effect that shifts the point of line intersection toward the first quadrant, thus mimicking a mixed inhibition pattern. The apparent *K*_*iu*_, which evidently emerges as a mere artifact, only depends on the [I]/[E] ratio: the smaller the ratio is, the smaller the *K*_*iu*_. As [I] tends to 0, however, the inhibition fades off, and the trace of the inhibited reaction tends to overlap with that of the uninhibited reaction. Therefore, under no circumstance, the intersection point can be below the 1/[S] axis, and hence, the apparent *K*_*iu*_ turns out to be necessarily greater than, or at best equal to, *K*_*ic*_.

**Figure 3.**
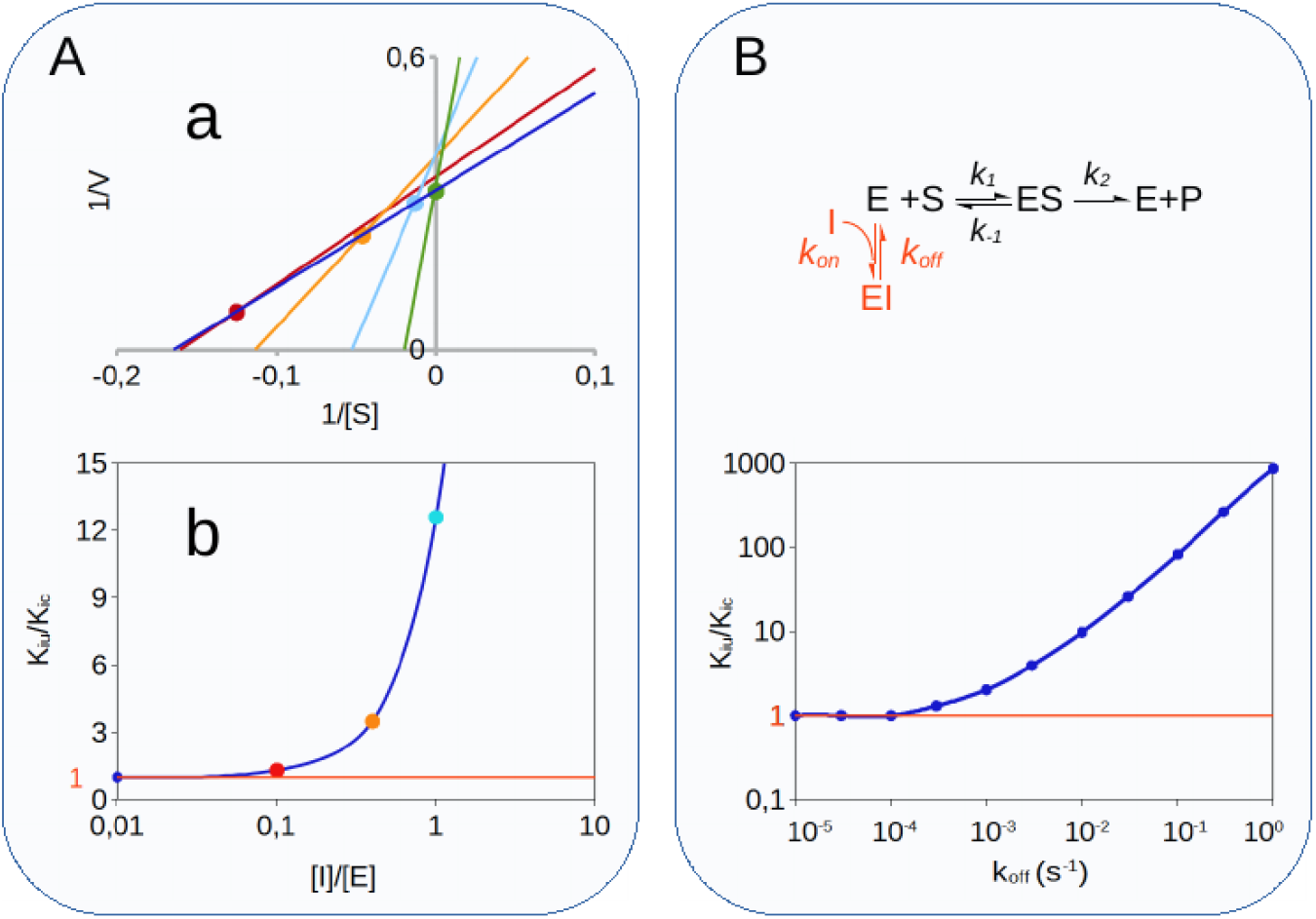
Simulation of MIM mechanisms of type 1. **Panel A**. Tight inhibition. (**a**) Double reciprocal plot of the uninhibited reaction (blue trace) and of reactions inhibited by four competitive inhibitors with different *K*_*i*_ and at different concentrations so that the [I]/*K*_*i*_ ratio is the same for all reactions. As expected for an active site-bound inhibitor, as long as the inhibitor concentration is at least ∼10 times larger than the enzyme concentration (for green trace [I]/[E]=10), the inhibition looks competitive. If [I] drops below 10·[E], the intercept shifts toward the first quadrant of the plot, mimicking a mixed-type inhibition whose *K*_*iu*_/*K*_*ic*_ ratio asymptotically tends to 1 as the inhibitor-to-enzyme ratio decreases (**b**). **Panel B**. Slow-dissociating inhibitors. For reactions initiated after a long enzyme-inhibitor incubation, the observed initial velocities depend critically on the dissociation rate constant of the inhibitor (*k*_*off*_). In the presence of active site-bound inhibitors, this mechanism generates an apparent mixed inhibition with *K*_*iu*_/*K*_*ic*,_ which asymptotically tends to 1 as the dissociation slows.

##### Case II – Time-dependent inhibition

The steady-state model assumes that the initial velocities are a direct measure of the steady-state rates. In the case of slow-binding inhibitors, however, the onset of the inhibition is delayed, as is the steady state, with the consequence that initial velocities underestimate the actual inhibition, leading to large overestimations of the *K*_*i*_s. Several methods to tackle slow-onset inhibition have been proposed [29–31], but because of their complexity, they have failed to turn into standard practices. A common strategy to tackle slow inhibition is instead to preincubate the enzyme and inhibitor and to add the substrate as the last reactant so that the enzyme-inhibitor equilibrium is attained before the reaction is started. In this case, if the enzyme-inhibitor dissociation is slow, the initial velocities underestimate the steady-state turnover, and the conventional steady-state analysis causes competitive inhibitors to be mistaken for mixed-type or noncompetitive. Although the effects of slow inhibitor dissociation on substrate-initiated reactions have long been described [32], the mechanism leading to the emergence of an apparent uncompetitive inhibition component has been elucidated only very recently [20]. As in all cases where the steady-state assumptions are not met, the output of the conventional data analysis is largely dependent on the experimental setup. In this specific case, the extent of the fictitious uncompetitive component strictly depends on the enzyme-inhibitor dissociation rate and on the dead time for the detection of initial velocities. Readers interested in deepening the mechanistic aspect of this kinetic artifact are referred to the cited literature [20]. Here, it will suffice to state that the apparent *K*_*iu*_ asymptotically tends to *K*_*ic*_ as the enzyme-inhibitor dissociation gets slower, eventually mimicking pure noncompetitive inhibition (*K*_*ic*_=*K*_*iu*_) when *k*_*off*_ is less than ∼10^−3^-10^−4^ s^-1^ (Figure 3B), so that necessarily *K*_*iu*_≥*K*_*ic*_.

#### 2.2.2. MIM mechanisms of type II – Complex non-Michaelian catalytic mechanisms

The Michaelis‒Menten model of enzyme kinetics relies on a simplified and incomplete approximation of the actual catalytic mechanisms [33]. If the role that these approximations play is not acknowledged, the interpretation of the analysis output can easily lead to incorrect conclusions.

##### Case III – Multisubstrate reactions

The Michaelis‒Menten model was derived for single substrate reactions, although these are quite rare in biochemistry and are confined to few isomerizations. It has been evaluated that some 60 percent of all the known enzyme-catalyzed reactions involve the conversion of two substrates into two products [34] according to the equation A+B=P+Q. For such reactions, referred to as Bi-Bi reactions, there are three common mechanisms: the ping-pong mechanism is observed in enzymes such as transaminases, which catalyze group transfer reactions where the transferred group relocates from substrate A to substrate B via an intermediate covalent enzyme adduct so that substrates bind one at a time in a compulsory order. In sequential mechanisms, on the contrary, substrates must occupy the enzyme active site at the same time. It can involve either a compulsory order of binding, as in most NAD-dependent dehydrogenases or CoA-dependent transferases where the coenzyme generally binds before the other substrate, or a random order where either of A or B can bind first. Depending on the actual mechanism, the steady-state analysis of competitive inhibitors of substrate A assayed against substrate B (i.e., at varying B concentrations and at fixed A) can give mixed-type, noncompetitive or uncompetitive inhibition. This is a very well-established fact thoroughly described in all enzymology manuals [35,36]. Here, we sought to demonstrate whether in the apparent mixed-type or noncompetitive inhibition observed for sequential mechanisms, the uncompetitive inhibition constant, 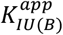, is necessarily always greater than or equal to the competitive constant, 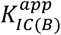.

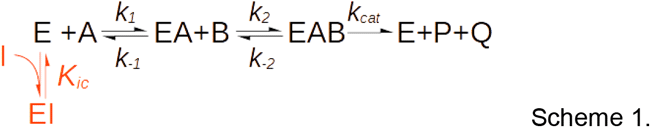

For a sequential reaction with compulsory order of binding, as depicted in Scheme 1, it can be shown (see Methods) that 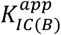 and 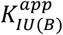 depend on the concentration of substrate A according to the following rules:

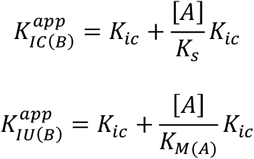

where *K*_*ic*_ is the true dissociation constant for the enzyme-inhibitor complex (i.e., the inhibition constant versus substrate A), *K*_*s*_ is the dissociation constant of the enzyme-substrate A complex and *K*_*M(A)*_ is the Michaelis constant for substrate A. At any given concentration of substrate A, the ratio between the apparent uncompetitive and competitive components is:

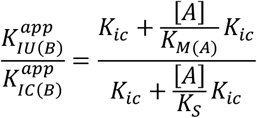

which asymptotically tends to 1 as the concentration of substrate A tends to 0 and to *K*_*s*_/*K*_*M(A*)_ as [A] tends to infinity. This 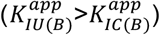 ratio dependence on [A] was confirmed by steady-state analysis of the KinTek Explorer simulations (Figure 4A). By definition, *K*_*s*_/*K*_*M(A*)_ is equal to *k*_*cat*_/*k*_*-1*_, which for a compulsory order mechanism is always greater than 1 (see Methods), hence leading to the conclusion that in this apparent mixed inhibition, the uncompetitive component is always necessarily greater than the competitive one 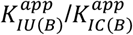. For sequential random-order mechanisms, the two inhibition components are identical, and it can be demonstrated that they are equal to:

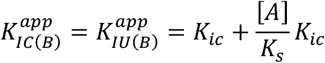

**Figure 4.**
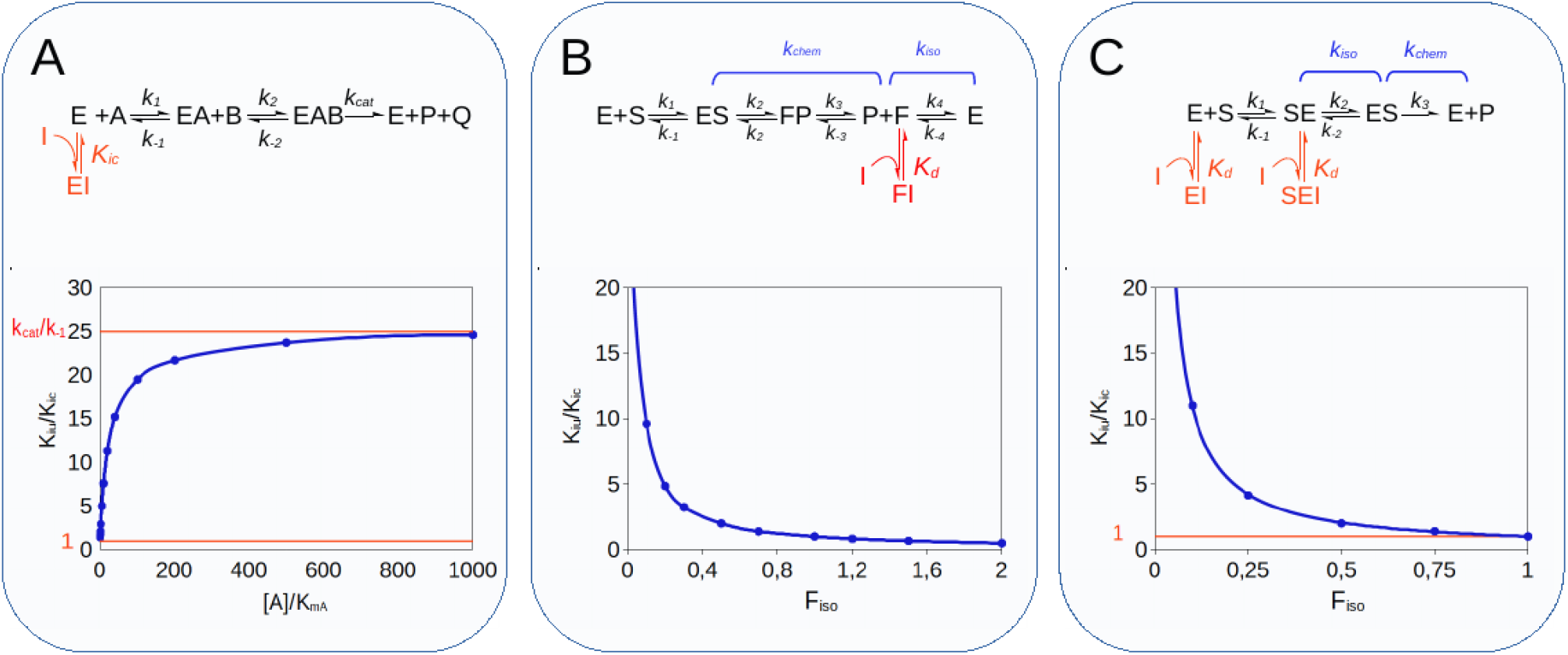
Simulation of MIM mechanisms of type 2. **Panel A**. Multisubstrate reactions. For bi-bi reactions with a sequential ordered mechanism, when the reaction is measured as a function of the substrate B concentration, competitive inhibitors directed against the binding site of substrate A generate an apparent mixed inhibition with a *K*_*iu*_/*K*_*ic*_ ratio that tends asymptotically to 1 when [A]/*k*_*mA*_ tends to 0 and to *k*_*cat*_/*k*_*-1*_ when [A] tends to ∞. **Panel B**. For iso-mechanism enzymes, inhibitors that target the product-binding isomer of the free enzyme (F in the reaction scheme) generate mixed-type inhibition with a *K*_*iu*_/*K*_*ic*_ ratio equal to 1/F_iso_. **Panel C**. With exo-site enzymes, active-site directed inhibitors cause mixed inhibition with *K*_*iu*_/*K*_*ic*_ that tends asymptotically to 1 as F_iso_ tends to 1.

Therefore, the 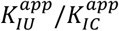 ratio is always equal to 1, and the apparent inhibition against substrate B always looks of the pure noncompetitive type.

According to the statistical study in chapter 2.1, approximately two-thirds of the enzymes for which mixed or noncompetitive inhibitors have been reported catalyze Bi-Bi reactions. This suggests that multisubstrate enzymes represent the most frequent source of apparent mixed-type inhibition.

##### Case IV – Iso-mechanisms

The cyclic nature of enzymatic catalysis implies that once the reaction products are released, the enzyme reverts to the form competent for the binding of the substrate. The Michaelis‒Menten model assumes that the isomerization between the substrate form and the product form of the enzyme is instantaneous and therefore can be neglected. This, indeed, is true in most cases. There are, however, exceptions, as isomerases that assume different conformations to bind different isomers, enzymes engaging in general acid or base catalysis that need reprotonation, or hydrolytic enzymes that require rehydratation [37,38]. If the isomerization step that restores the initial enzyme form is slow enough to affect catalytic turnover, the enzyme is said to have an isomechanism (scheme 2) [39].

Compelling analytical tools to investigate iso-mechanisms were developed only in the 1990s, mainly by Dexter Northorp, who exploited the distinctive inhibition pattern of these enzymes. In the absence of an isomechanism, the reaction product is typically a competitive inhibitor of the substrate because both reactants bind the same form of enzyme. In contrast, for enzymes with iso-mechanisms, substrate and product bind two distinct enzyme forms; therefore, the product inhibition pattern is of the mixed-type [40,37]. Analogous considerations apply to the case of dead-end inhibitors that target exclusively the product form of the active site.

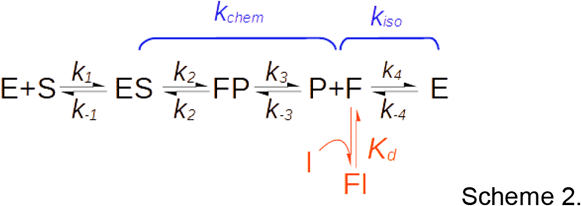

As shown in scheme 2, for iso-mechanisms, the overall turnover rate, (*k*_*cat*_), is made up of two segments, the catalytic step governed by the net rate constant *k*_*chem*_, and the isomerization step, with net rate constant k_iso_. The binding of an inhibitor to the product form of the enzyme (F in Scheme 2) causes a mixed-type inhibition with:

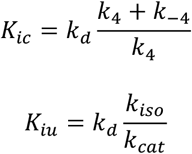

with *K*_*d*_ representing the dissociation constant of the FI complex (see Methods). As correctly predicted by Northorp [37–39], under the initial velocities condition, the *K*_*ic*_/*K*_*iu*_ ratio equals F_iso_ (Figure 4B), a parameter that quantifies the kinetic significance of the isomerization segment and that is defined as:

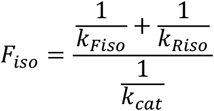

where *k*_*Fiso*_ and *k*_*Riso*_ are the rate constants for the forward and reverse isomerization steps, respectively.

Unlike the previous MIM mechanisms, in this case, the reduction in the turnover rate caused by active site-bound inhibitors is not just an artifact. Because F_iso_ depicts an intrinsic feature of the catalytic mechanism, all inhibitors of a given enzyme would cause a mixed inhibition with an identical *K*_*iu*_/*K*_*ic*_ ratio. Unfortunately, there are no reliable data on the prevalence of iso-mechanisms, so it is difficult to estimate their contribution to the overall occurrence of mixed inhibition. It is worth noting, however, that the product form of the free enzyme is a short-lived molecular species generated along with the catalysis rather than a stable enzyme isoform. This implies that isomerization in the forward direction (F→E) is necessarily faster than in the reverse direction (F←E) [41], which is consistent with the finding that the chemical segment is always the rate-limiting step [37]. Consequently, even though the *K*_*iu*_s generated by iso-mechanisms cannot be algebraically shown to be necessarily greater than the *K*_*ic*_s, mechanistic considerations explain why F_iso_ is practically always less than 1 and, consequently, why, in this case too, the mixed inhibition is dominated by the competitive component.

##### Case V – Exo-sites

With most enzymes, it is not possible to operate a clear distinction between the portion of the active site involved with the binding of the substrate and the actual catalytic site because the catalytic site also contributes to substrate stabilization, at least to some extent. In this respect, all enzymes catalyzing the covalent modification of large macromolecular substrates, endonucleases [42], protein kinases/phosphatases [43], proteases [45], etc., represent a notable exception. Often, this type of enzyme derives most of the affinity for its substrate from the region of the protein surface that is distinct and remote from the catalytic site. The substrate recognition site, in this case, is called an exo-site [45].

For exo-site enzymes, the binding of the substrate typically proceeds through two sequential steps (Scheme 3), the first leading to an “encounter complex”, here denoted as “SE”, with the substrate bound only to the exosite. SE then decays monomolecularly to form the catalytically competent complex “ES”, with the consensus sequence of the substrate accommodated within the active site and ready to undergo catalytic conversion.

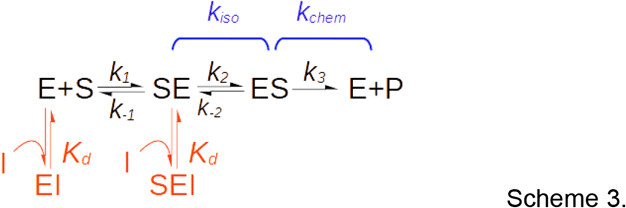

An active site-directed inhibitor, I, can bind both the free enzyme E and the encounter complex SE but not ES. Or in other words, the inhibitor does not compete with the initial enzyme-substrate recognition but impedes the isomerization of the SE complex that leads to the formation of the catalytically competent complex ES. This causes a reduction of the “degree of competition” that in a double reciprocal plot is visualized as a mixed-type inhibition pattern [13].

To model this reaction mechanism, it is convenient to regard *k*_*cat*_ as composed of two segments, the isomerization of the SE complex and the chemical catalytic step, governed by the constant rates *k*_*iso*_ and *k*_*chem*_, respectively (Scheme 3).

Analogously to the previous iso-mechanism case, the coefficient F_iso_, here defined as *k*_*cat*_/*k*_*iso*_, quantifies the kinetic relevance of the isomerization step. F_iso_ assumes values between 0 when *k*_*iso*_ is so large that the isomerization is kinetically irrelevant and 1 when it is slow enough to be completely rate-limiting.

Simulation of this reaction with KinTek Explorer (Figure 4C) showed that the *K*_*iu*_/*K*_*ic*_ ratio is equal to 1/F_iso_. From which follows that if F_iso_ tends to 0, the inhibition is competitive, as expected for an active site-bound inhibitor. If F_iso_=1, the catalytic segment of the reaction can be neglected, and the inhibition turns pure noncompetitive (*K*_*ic*_=*K*_*iu*_). For 0<F_kiso_<1, the inhibition is of the mixed-type with *K*_*ic*_<*K*_*iu*_, so that necessarily *K*_*iu*_≥*K*_*ic*_ (see Methods).

The reduction of the affinity for the substrate and of the apparent maximal velocity are both actual. However, as for the previous iso-mechanism case, and contrary to what the conventional mixed inhibition theory mandates, there is no allotopic site involved, and *K*_*iu*_ is not a true dissociation constant but just a coefficient that relates the extent of *V*_*max*_ reduction to the reduction of the apparent *k*_*M*_*/V*_*max*_.

Although enzymes with exosites are often of utmost pharmacological relevance [13], the proper study of their inhibition mechanism, which necessarily has to be performed using the physiologic substrates, has been limited by the remarkable effort required to produce enough of the proteinaceous substrates and to measure the time course of their chemical modification. Indeed, this type of study has been reported for just a handful of enzymes, mostly serine proteases of the coagulation cascade process [45–47], so that experimental evidence is scant and the contribution of exo-site enzyme inhibition to the statistics of chapter 2.1 is practically null.

## 3. Discussion

The notion of mixed and noncompetitive inhibition being caused by inhibitor binding to two distinct sites lays at the foundation of the current enzyme theory [48] and is deeply rooted in most biochemists. Several authors agree that enzyme inhibition should be regarded as always of the mixed type, with pure competitive or uncompetitive inhibition being the limit cases occurring when either *K*_*iu*_>>*K*_*ic*_ or *K*_*ic*_>>*K*_*iu*_, respectively [9,49]. Although from a purely mathematical perspective this might still be correct, the considerations presented herein suggest that the nature of enzyme inhibition is almost always competitive, with rare examples of true uncompetitive inhibition, probably accounting for less than 5% of total cases, and with mixed-type inhibition being substantially insignificant.

It has to be pointed out that already in 2004 Cornish-Boweden suggested caution in the interpretation of mixed inhibition patterns [9]. However, perhaps because his considerations were not supported by rigorous analytical demonstrations, it does not seem to have contributed appreciably to the general understanding of mixed inhibition and, as a matter of fact, the classic two-site model has continued to be perpetuated through countless scientific articles and enzymology textbooks (see as notable examples references [4] and [13]).

Given the huge diversity displayed by enzymes, in terms of both structures and mechanisms, the possibility of mixed inhibition occurring in the same form as envisaged in the Michaelian model cannot be positively ruled out. Yet, the finding that the five MIM mechanisms generate apparent mixed inhibition with *K*_*iu*_s that are always larger than their cognate *K*_*ic*_s constitutes a statistical evidence of sufficient force to prove that the inhibitors reported to be of the mixed type, practically none of which binds preferentially the allotopic sites, are, with virtually no exception, just miss evaluated cases of active site-bound inhibitors.

It is worth emphasizing that, far from being of a mere academic interest, this “competitive versus mixed-type” issue heavily impacts our understanding of the actual biological and pharmacological role played by reversible enzyme inhibitors. One immediate *in vivo* effect of an enzyme’s inhibition, in fact, is the increase in its substrate(s) concentration. As shown earlier [10,50], the dependence of substrate accumulation on the competitive inhibitor concentration in the first approximation can be regarded as linear. Conversely, increasing the dose of a mixed (or uncompetitive) inhibitor causes an exponential-like increase in the substrate, which can hardly be tolerated by the cell. This implies two consequences: first, the mechanism of inhibition is critical to predict the pharmacodynamic properties of a drug [51,52], and hence, it should be regarded as a key factor in the development of guidelines for pharmacological research [13,53]. Second, uncompetitive and mixed inhibition by naturally occurring metabolites is so detrimental to living organisms that in nature they have to be very rare. A prediction, this latter, which has been for a long time contradicted by the reported common occurrence of mixed inhibition but that finds corroboration in the evidence discussed herein.

## 4. Conclusion

The outcome of this theoretical investigation is that the accepted two-site theory of mixed enzyme inhibition is a blunder. Three out of the five mechanisms that mimic mixed inhibition are merely a consequence of inattentive evaluation of experimental conditions (cases I and II) or of misinterpretation of the results (case III). In these three cases, the inhibition is purely competitive, and the observed uncompetitive component is just an artifact deprived of any biological significance.

In principle, cases IV and V can actually result in a concomitant reduction in substrate affinity and maximal velocity. However, it must be stressed that they are both very uncommon. Iso-mechanisms (case IV) have been reported only for a few enzyme types and occur primarily as a form of product inhibition [9]. With exosite enzymes (case V), mixed inhibition can only be observed if the enzyme activity is measured against physiologic macromolecular substrates. However, this kind of experiment, due to the high cost, has been carried out only very rarely and just for very few enzymes.

## 5. Methods

### 5.1. Simulation of mixed inhibition-mimicking mechanisms

To simulate MIM mechanisms, initial velocities were generated using KinTek Explorer, and the inhibition constants *K*_*ic*_ and *K*_*iu*_ were determined by the Dixon and Cornish-Bowden plots, respectively. Velocities were detected 60 seconds after the start of the reactions. For each mechanism, the reaction scheme, kinetic constants, and the concentration of reagents are given below.

#### Case I – Tight binding inhibitors

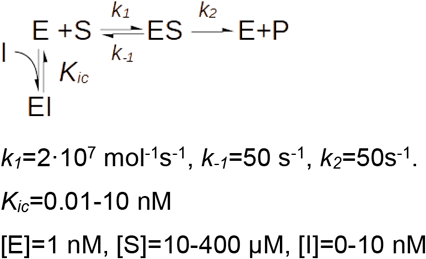

#### Case III – Multisubstrate reactions

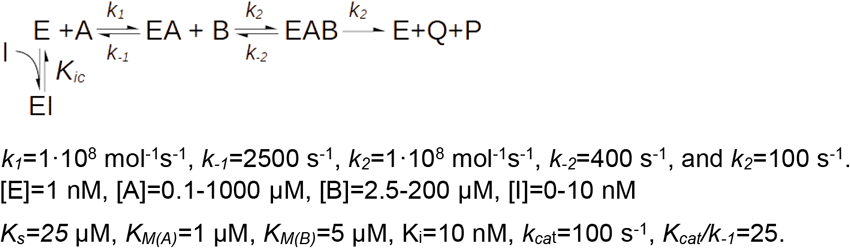

#### Case IV – Iso-mechanisms

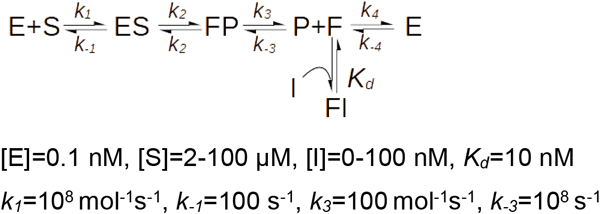

| F<sub>iso</sub> | 0 | 0.1 | 0.2 | 0.5 | 0.7 | 1 |
| --- | --- | --- | --- | --- | --- | --- |
| $k_2 \text{ (mol}^{-1}\text{s}^{-1}\text{)}$ | 100 | 111 | 125 | 200 | 333 | $10^6$ |
| $k_{-2} \text{ (s}^{-1}\text{)}$ | 100 | 111 | 125 | 200 | 333 | $10^6$ |
| $k_4 \text{ (mol}^{-1}\text{s}^{-1}\text{)}$ | $10^6$ | 1000 | 500 | 200 | 143 | 100 |
| $k_{-4} \text{ (s}^{-1}\text{)}$ | $10^6$ | 1000 | 500 | 200 | 143 | 100 |

#### Case V – Exo-site

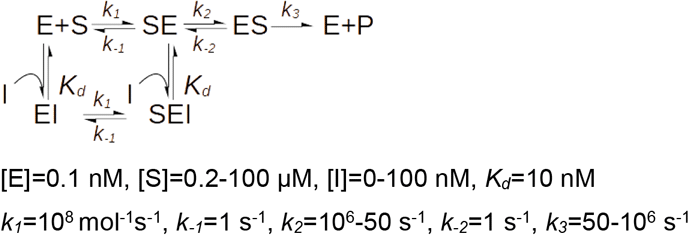

### 5.2. Derivation of the *Kiu*-*Kic* relation

#### Case III – Multisubstrate reaction

##### Sequential reaction with compulsory order

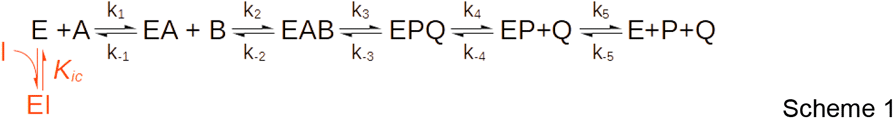

For a Bi-Bi reaction with a sequential ordered mechanism as depicted in Scheme 1, the equation for the initial rate is:

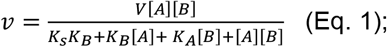

where *V* is the limiting reaction velocity at saturating A and B substrate concentrations, *K*_*A*_ and *K*_*B*_ are the Michaelis constants for A and B, respectively, and *K*_*S*_ is the dissociation constant for substrate A, equal to *k*_*-1*_*/k*_*1*_. At a fixed substrate A concentration, the apparent maximal velocity 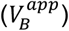 observed at saturating B and the apparent Michaelis constant for substrate B 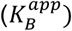 are both a function of the concentration of A:

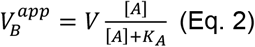

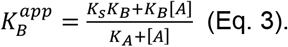

In the presence of binding site A-directed inhibitor I, the apparent Michaelian parameters are:

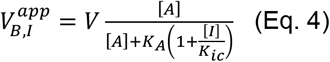

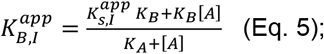

where 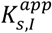 is the apparent substrate A dissociation constant observed in the presence of I: 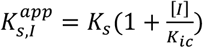.

By definition, mixed inhibition affects both the *V/K*_*M*_ ratio and *V*, so that:

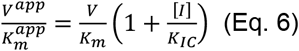

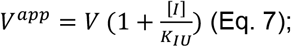

where *K*_*IC*_ and *K*_*IU*_ are the true competitive and uncompetitive inhibition constants, respectively.

To derive the relation between the true dissociation constant of inhibitor I (*K*_*ic*_) and the apparent *K*_*ic*_ and *K*_*iu*_ (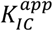 and 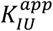) observed when the inhibitor is assayed at constant A against substrate B, the equations for the apparent *V/V*^*app*^ and *V*/*K*_*M*_ */(V*/*K*_*M*_)^*app*^ (i.e., Eq.6 and Eq.7) were equated to Eq.4/2 and Eq.5/3.

###### Derivation of the K_ic_-K_IC_^app^ relation

For the reaction of Scheme 1, at constant A and variable B, *V/K*_*m*_ is:

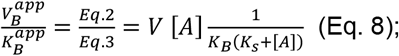

and, in the presence of the competitive inhibitor I, *(V/K*_*M*_)^*app*^ is:

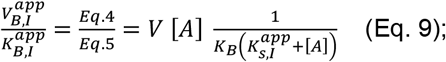

Therefore, the ratio between *V/K*_*M*_ and *(V/K*_*M*_)^*app*^ is:

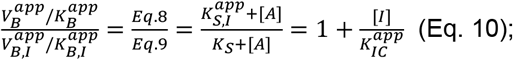

where 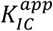 represents the apparent competitive inhibition constant against substrate B caused at a fixed A concentration by the binding site A–directed inhibitor I.

From equation 10 follows that:

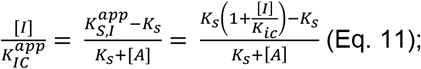

and hence that:

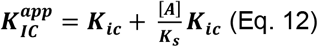

###### Derivation of the K_ic_-K_IU_^app^ relation

For the reaction of Scheme 1, at constant A and variable B, *V* is 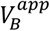 (see Eq. 2), and in the presence of inhibitor I, *V*^*app*^ is 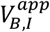 (see Eq.

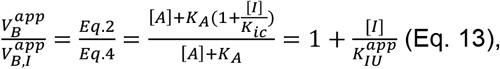

where 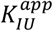 is the apparent uncompetitive inhibition constant against substrate B caused at a fixed A concentration by the binding site A – directed inhibitor I. From equation 13 follows that:

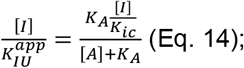

Hence,

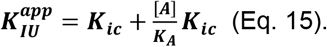

In conclusion, binding site A-directed inhibitor I causes an apparent mixed-type inhibition against substrate B characterized by a *K*_*IU*_*/K*_*IC*_ ratio that is:

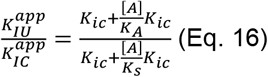

Because 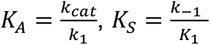 and 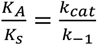, it follows that:

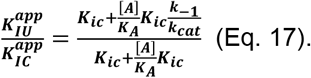

From equation 17, it is evident that 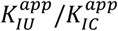 tends asymptotically to 1 if [A] tends to 0 and to 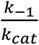 if [A] tends to infinity. This implies that 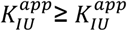 if *k*_*-1*_ ≥ *k*_*cat*_.

Mechanistic considerations suggest that *k*_*-1*_ is necessarily larger than *k*_*cat*_:

the overall turnover rate, *k*_*cat*_, results from the combination of the constant rate for the actual catalytic conversion of substrate into products (*k*_*3*_) and the net rates for the dissociation of the products P and Q (see Scheme 1), according to the equation:

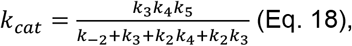

from which it is evident that *k*_*cat*_ is necessarily smaller than any of *k*_*3*_, *k*_*4*_ or *k*_*5*_, and specifically that *k*_*5*_>>*k*_*cat*_.

In enzymes exhibiting a compulsory order mechanism, only the binding site of substrate A is constitutively present on the free enzyme. The binding of A then induces a structural rearrangement that results in the formation of the binding site for substrate B, which explains why B can only bind after A. k_-1_ is the kinetic constant for the reverse of this process, that is, the dissociation of the EA complex and the consequent reversion to the constitutive free enzyme form. This transition is identical to that occurring at the very end of the catalytic turnover, when the product Q is released and the enzyme reverts to the free form E. Hence, because the two microscopic constants *k*_*-1*_ and *k*_*5*_ describe the same process, whose rate limiting step is the protein structural rearrangement, they can be expected to have similar values. Moreover, it should be considered that the rate-limiting step of enzymatic turnover often is the chemical conversion, in this case represented by the constant *k*_*3*_. So that, in conclusion, since *k*_*5*_>>*k*_*cat*_ and *k*_*-1*_≃*k*_*5*_, follows that *k*_*-1*_>*k*_*cat*_, and hence, 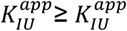.

##### Sequential reaction with random order

For sequential random-order reactions, the initial velocity equation is:

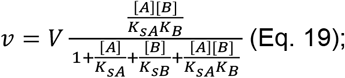

where *K*_*sA*_ and *K*_*sB*_ are the dissociation constants for substrates A and B, respectively.

At a fixed A concentration, the apparent maximal velocity measured at saturating B and the apparent Michaelis constant for B are:

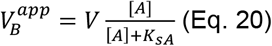

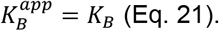

In the presence of the substrate A binding site-directed inhibitor I, the apparent Michaelian parameters are:

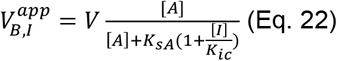

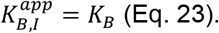

###### Derivation of the K_ic_-K_IC_^app^ relation

For the reaction of Scheme 1, at constant A and variable B, *V/K*_*m*_ is:

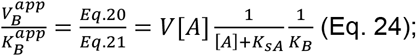

The *V*^*app*^*/K*_*m*_^*app*^ ratio caused by substrate A binding site-directed inhibitor I is:

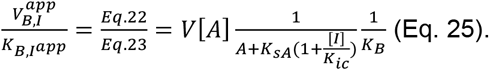

Thus, the ratio between *V/K*_*m*_ and *(V/K*_*m*_)^*app*^ is:

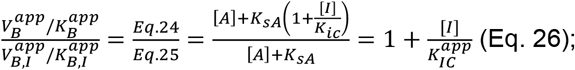

where 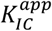 represents the apparent competitive inhibition constant against substrate B caused at a fixed A concentration by binding site A – directed inhibitor I.

From equation 26 follows that:

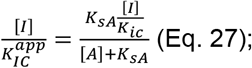

Hence,

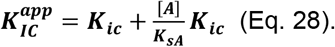

###### Derivation of the K_ic_-K_IU_^app^ relation

For the reaction of Scheme 1, at constant A and variable B, *V* is 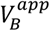 (see Eq. 24), and in the presence of inhibitor I, *V*^*app*^ is 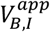 (see Eq. 25), so that the *V/V*^*app*^ ratio is given by:

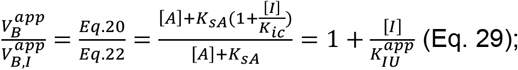

where 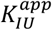 represents the apparent uncompetitive inhibition constant against substrate B caused at a fixed A concentration by binding site A – directed inhibitor I.

From equation 29, it follows that:

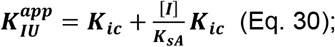

Hence, for sequential reactions with random order, 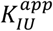 is always equal to 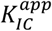, and consequently, the inhibition results are necessarily pure noncompetitive.

#### Case IV – Iso-mechanism

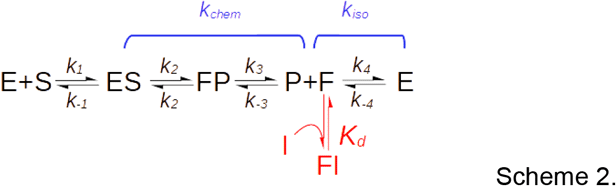

In iso-mechanism reactions, the turnover rate is made up by the contribution of two components, the chemical segment, which accounts for the actual catalytic step and for the release of the reaction product, with net rate *k*_*chem*_, and the isomerization segment, which accounts for the conversion of the enzyme product form (F) into the initial substrate form (E), with net rate *k*_*iso*_. The overall turnover rate, *k*_*cat*_, is given by:

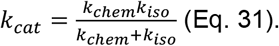

The inhibitor “I”, which binds the F form of the enzyme with a dissociation constant equal to *K*_*d*_, does not affect *k*_*chem*_ but reduces the apparent *k*_*iso*_ according to:

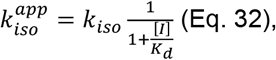

Consequently, the apparent *k*_*cat*_ observed in the presence of I is:

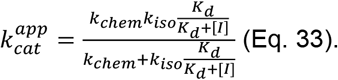

Due to the presence of a fast equilibrium between E and F, the binding of I to F causes a competitive inhibition whose *K*_*ic*_ depends both on the *K*_*d*_ and the forward and reverse isomerization rates:

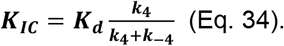

By definition, for uncompetitive inhibition, we have that:

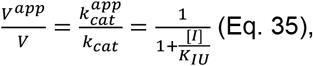

where *K*_*IU*_ is the uncompetitive inhibition constant. To derive the relation between the true dissociation constant (*K*_*d*_) and the observed uncompetitive inhibition constant (*K*_*IU*_), 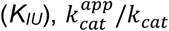 (Eq. 33/Eq. 31) ratio was equated to 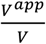 (Eq. 35):

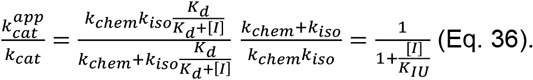

Form equation 36, by isolating the *K*_*iu*_ and *K*_*d*_ terms, follows that under the initial velocity assumption:

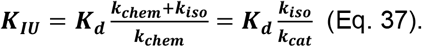

#### Case V – Exo-site

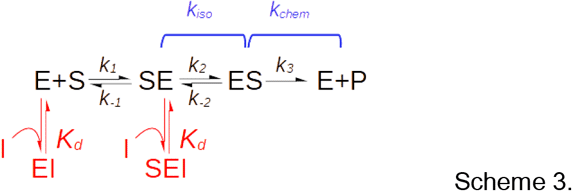

Scheme 3 depicts the mechanism of an exo-site enzyme. Similar to the previous Iso-mehanism case, the turnover rate can be regarded as made up of two contributions, the isomerization of the encounter complex (SE) to form the catalytically competent complex (ES), with net rate *k*_*iso*_, and the actual catalytic step, with net rate *k*_*chem*_, so that the overall turnover rate (*k*_*cat*_) is given by:

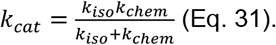

If we assume that the exo-site and active site are completely independent and hence that the interaction of the substrate with the exo-site does not affect the binding of the inhibitor, the *K*_*ic*_ determined by steady-state analysis corresponds to the actual EI (or SEI) dissociation constant (*K*_*d*_) independent of the kinetic relevance of the isomerization step. Hence:

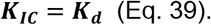

The binding of the active-site directed inhibitor I to the ES complex affects the apparent *k*_*iso*_ so that:

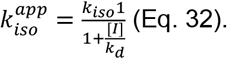

Consequently, the apparent *k*_*cat*_ observed in the presence of I is:

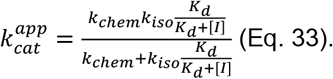

From which follows (see previous demonstration) that:

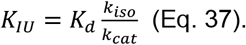

The *K*_*iu*_/*K*_*ic*_ ratio can be obtained by simply combining equations 39 and 37:

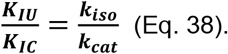

